# Quantifying and predicting chromatic thresholds

**DOI:** 10.1101/2023.06.06.543898

**Authors:** Jingyang Zhou

## Abstract

Perceptual thresholds measured in the two-dimensional chromatic diagram are elliptical in shape. Across different parts of the chromatic diagram, these ellipses vary in their sizes, their tilting angles, and in how much they elongate. Overall, the chromatic thresholds exhibit intriguing patterns that were reflected in McAdam’s measurements in 1942. Previously, da Fonseca and Samengo (2016) used a neural model combined with Fisher information (a quantification of perceptual thresholds) to predict the pattern of chromatic thresholds measured in human observers. The model assumes linear cone responses paired with Poisson noise. I furthered the analysis, and studied two additional aspects of chromatic perception. First, I quantified how the pattern of chromatic thresholds vary when the proportion of three cone types (short-, mid-, and long-wavelength) varies. This analysis potentially leads to efficient estimation of thresholds across the chromatic diagram. Second, I analyzed to what extent the assumption of Poisson noise contributes to the threshold predictions. Surprisingly, eliminating Poisson noise betters the model prediction. So in addition to Poisson noise, I examined three alternative noise assumptions, and achieved improved predictions to MacAdam’s data. At last, I examined an application using the improved model-predictions. The total number of cones, as well as the proportion of *S* cone vary across retinal eccentricities. I showed that these two variations predict chromatic threshold patterns across retinal eccentricities are drastically different.

## 1 Introduction

Perceptual discriminability relates an observer’s external performance (e.g. observers’ “yes” and “no” answers in two-alternative forced choices) to internal stimulus representations (e.g. Green and Swets 1966; F. A. A. Kingdom and Prins 2009). Signal detection theory assumes these internal representations are stochastic, and the stochasticity gives rise to observers’ often inconsistent choices when perceptual tasks become difficult (e.g. discriminating between two near-identical images). Suppose a stimulus *s* varies along one dimension, and an observer’s internal representation is modeled as a one-dimensional distribution *p*(*r*| *s*). Many empirical studies (e.g. Green and Swets 1966; F. A. A. Kingdom and Prins 2009; Geisler 2002) assumes *p*(*r*| *s*) is Gaussian-distributed, with the Gaussian mean *µ*(*s*) varies with the stimulus level *s*, and other aspects of the representation, such as variance *σ*(*s*)^2^, stay fixed for all stimuli.

To relate perceptual discriminability to internal representations, one can compute Fisher information of *p*(*r*| *s*). (Stein, Mezghani, and Nossek 2014; J. Zhou, Duong, and E. P. Simoncelli 2022; Seung and Sompolinsky 1993; May and J. Solomon 2015; Series, Stocker, and E. P. Simoncelli 2009; Kanitscheider, Coen-Cagli, and Pouget 2015; Kfashan et al. 2021; Brunel and Nadal 1998; Abbot and Dayan 1999). Fisher information is a statistical tool that quantifies how much a distribution varies when its parameters change. In our case, we can compute how much the internal stimulus representation *p*(*r* |*s*) changes, when *s* is adjusted slightly. Square-root of the Fisher information — *Fisher discriminability*, summarizes perceptual discriminability *d*(*s*) as the following:

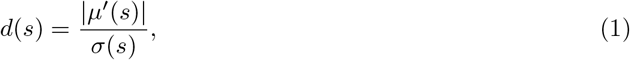

and *µ*^′^(*s*) is the derivative of the representational mean, and *σ*(*s*) is the representation standard deviation, which may also vary with stimulus levels (e.g. J. Zhou, Duong, and E. P. Simoncelli 2022).

Perceptual discriminability can also be studied by varying along more than a single stimulus dimension at a time. Fisher information naturally generalizes to quantifying perceptual discriminability when an image is perturbed in multiple dimensions (e.g. Berardino et al. 2018). For example, when an arbitrary color becomes slightly more red, slightly more green, or slightly more blue, the extent of perceptual change that we experience can be quite different (MacAdam 1942; Poirson and Wandell 1990; Brandt and Vorobyev 1997). Combined with a neural model, Fisher information can be used to predict perceptual discriminability along all three color perturbation directions, as well as along any combination of the three directions (Fonseca and Semengo 2016). In general, suppose a *p*-dimensional stimulus **s** is encoded by *n* number of neurons’ stochastic responses. I use ***µ***(**s**), an *n*-dimensional column vector, to denote the neurons’ mean response rates to stimulus **s**, and I use Σ(**s**), the *n* × *n* covariance matrix, to account for co-variation between neurons when they respond to repeated presentations of the same stimulus. To account for perceptual discriminability to arbitrary stimulus perturbations to **s**, we can compute the lower-bound on Fisher information (Stein, Mezghani, and Nossek 2014; J. Zhou, Duong, and E. P. Simoncelli 2022), which has an expression closely related to Fisher discriminability for one-dimensional stimulus perturbation (Equation 1):

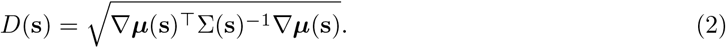

∇***µ***(**s**) is the Jacobian of neuronal response ***µ***(**s**) with respect to stimulus. Using this multi-dimensional Fisher expression, in this paper, I examined perceptual thresholds – the inverse of perceptual discriminiability, in chromatic diagram (2-dimensional color space with fixed luminance).

MacAdam measured two-dimensional perceptual thresholds in the chromatic diagram (MacAdam 1942), and the thresholds exhibit intricate patterns across the chromatic diagram. At a single color (a reference color), perceptual thresholds measured by perturbing the color along different directions exhibit an elliptical shape around the reference color (Figure 1A). Threshold measured at a reference color is the largest along the major axis of the ellipse, and the smallest along the minor axis (notice that originally, MacAdam’s 1942 experiment was done using a matching procedure). At different reference colors, the elliptical representations of chromatic thresholds change in shape, size, and how much they tilt. Existing literature used a linear neural model paired with Poisson noise to quantify chromatic thresholds (or “MacAdam ellipses”), and explained large extent of data variance (Fonseca and Semengo 2016). In this paper, I first re-produced this model. Then, I varied the proportion of each cone type (short-, mid-, and long-wavelength) and examined how chromatic thresholds correspondingly vary. This analysis potentially leads to measuring thresholds across the chromatic diagram with much improved efficiency. I further improved model-predicted chromatic thresholds by pairing the linear neural model with different types of plausible neuronal noise. At the end, I showed an example of using model-predicted chromatic thresholds to study how our color vision differs across retinal eccentricities.

**Figure 1:**
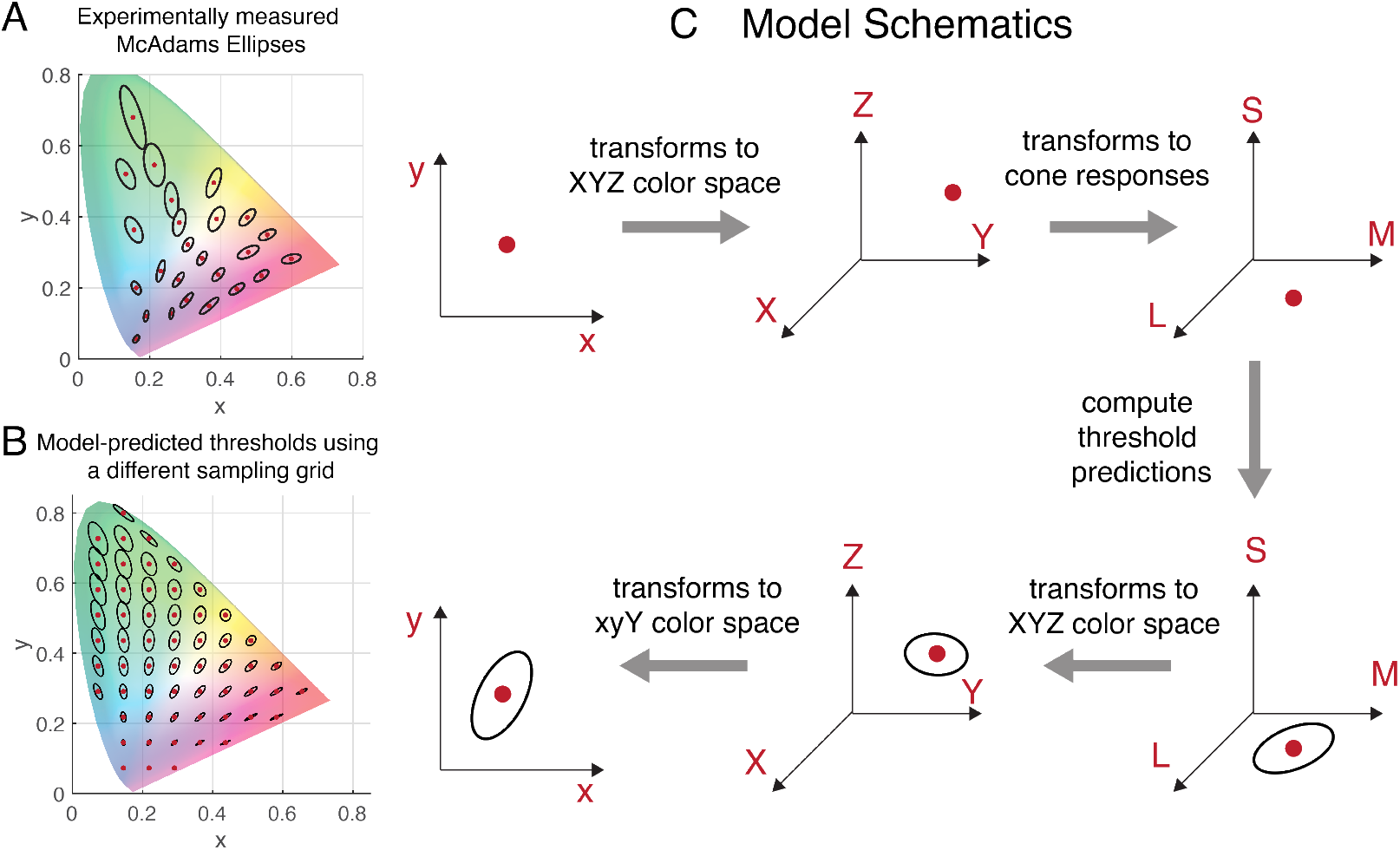
Model schematics. A. MacAdam ellipses. B. Predicted chromatic thresholds using a linear neural model paired with Poisson noise. Reference colors were sampled on a regularly spaced grid on the chromatic diagram. C. Model schematics. First, I sampled reference colors in the 2D chromatic space (the *xy* space). Then I mapped these sampled reference colors to the *XY Z* color space, from which I further projected the sampled colors to the space of cone responses (the LMS space). To visualize predictions of chromatic thresholds, I generated threshold predictions in the LMS space by computing *Fisher thresholds* – the inverse of Fisher discriminability. I mapped these threshold predictions back to the *XY Z* space, then further back to the *xy* space.

## 2 Results

### 2.1 Predicting chromatic thresholds using a linear neural model paired with Poisson noise

The process of predicting chromatic thresholds (Fonseca and Semengo 2016) can be partitioned into two steps. In the first step, I sampled reference colors in the *xy* chromatic diagram, and mapped them to the space of cone responses. Reference colors in the *xy* space could be sampled on a regularly spaced grid as in Figure 1B, or it could be more irregularly sampled, as in the case of the MacAdam ellipses. Mapping these samples to the space of cone responses takes two transforms (Figure 1C). First, I mapped these samples to the *XY Z* color space (via a non-linear function). Then via a linear transform, I further mapped the samples to the space of cone responses (the LMS space) (Lu and Dosher 2014; Fairman, Brill, and Hemmendinger 1997).

As the second step of generating chromatic threshold predictions, I derived thresholds using cone responses by computing the inverse of Fisher discriminability (i.e. *Fisher thresholds*). To visualize thresholds in the chromatic diagram, I transformed the Fisher thresholds back to the *xy* space. Here are the detailed computation.

### Forward direction: From *xy* to neural response space

The *xy* chromaticity, or the *xyY* color space, assumes a fixed luminance level *Y*, the value of which I empirically estimated from MacAdam’s data. Notice that previous evidence suggested that the shapes of the MacAdam ellipses do not vary much for a range of luminance (Brown and MacAdam 1949). To map the sampled reference colors from the *xyY* space to the space of cone responses takes two transforms. First, I mapped the reference colors from the *xyY* space to the *XY Z* space via a non-linear transform. To be specific, I use **c** to indicate a reference color sampled in the *xyY* space, and **c** is a three-dimensional column vector **c** = [*x, y, Y*]^T^. The first two entries of **c** indicate its chromatic coordinate, and *Y* – the luminance level, was empirically estimated. To map **c** to the *XY Z* space, I first examined the non-linear transform from *XY Z* to *xyY* space (because it is intuitively defined), then I computed the inverse of this transform. The non-linear transform 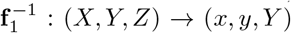 separates luminance and chromaticity, and has the following expression:

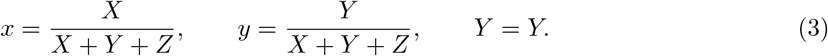

The inverse of this map, **f**_1_ : (*x, y, Y*) → (*X, Y, Z*), can be expressed as:

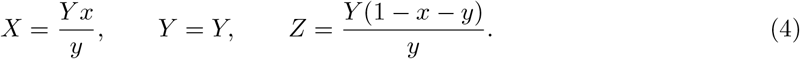

In the second step, I projected the reference colors from the *XY Z* space to the *LMS* cone response space through a linear transform. I used ***α*** = [*α*_*S*_, *α*_*M*_, *α*_*L*_]^⊤^ to indicate the averaged cone response rate. In this paper, ***α*** can be viewed as the stimulus input to the model. Following the notation convention in Fonseca and Semengo 2016, I used ***β*** = [*β*_*S*_, *β*_*M*_, *β*_*L*_] to indicate the proportion of *S, M* and *L* cones in a retina, and *β*_*S*_ + *β*_*M*_ + *β*_*L*_ = 1. The transform from *XY Z* color space to the *LMS* space is linear, so it can be represented using a 3 ×3 matrix *F*_2_. In Fonseca and Semengo 2016, *F*_2_ was empirically estimated as:

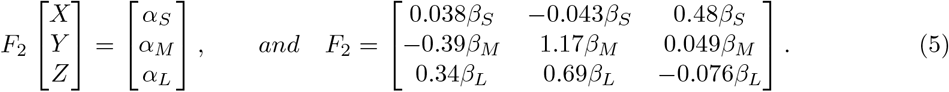

Notice that in the above analysis, both transforms **f**_1_ and *F*_2_ are invertible. Now I use the following equation to summarize the transform in the forward direction:

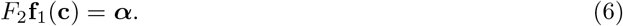

### Backward direction: from neural response space back to *xy* space

For the backward direction, I first derived perceptual thresholds by computing Fisher discriminability using neural response ***α***. Then, I mapped the predicted thresholds back to the *xyY* chromatic space.

To compute Fisher discriminability using Equation 2, I need to know the mean and the covariance of the neural responses. In the linear response model, the mean neural response is assumed to coincide with the three-dimensional light input ***α***, and the derivative of the mean response (with respect to ***α***) is the identity matrix **I**. The model assumes Poisson noise, and Poisson noise implies that each neuron’s response variance is identical to the mean. For example, the response variance for *S* - cones, 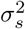, is identical to the mean response, and 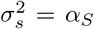. Using Equation 2, I found the following expression for Fisher discriminability *D*(***α***), as well as Fisher thresholds, which is the matrix inverse of discriminability:

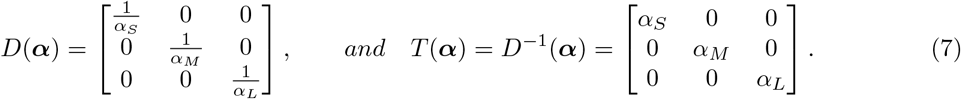

To visualize thresholds in the *xyY* space, I mapped *T* (***α***) to the *xyY* space in two steps. First, we can make an observation that to transform neural response ***α*** back to the *xyY* space, we need to compute 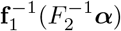 To transform thresholds *T* (***α***) – a quadratic form, back to the stimulus space, I computed the following (see Methods for details):

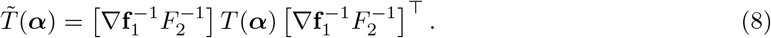

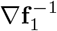is the Jacobian of the function 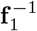 with respect to input 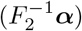, and 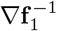 has the following form:

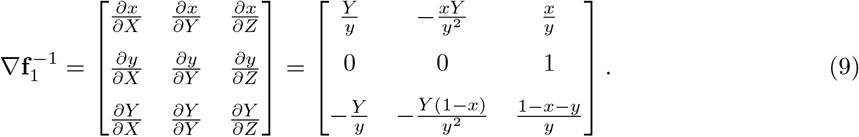

There is one additional transform in computing 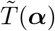, the Jacobian of map 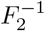. Because 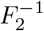 is a linear map, 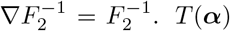 describes a three-dimensional ellipsoid centered at ***α*** in the *LMS* space, and after the two Jacobian transforms, 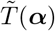 describes a three-dimensional ellipsoid in the *xyY* space. To visualize perceptual thresholds in the two-dimensional *xy* space, as in Fonseca and Semengo 2016, I extracted the 2 ×2 sub-matrix of 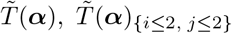, that represents the metric tensor restricted to the *xy* plane. In Figure 1C, I illustrated both the forward and the backward direction of computing model-predicted chromatic thresholds.

### 2.2 Efficiently estimating chromatic thresholds

Using the cone model described above, I studied two questions in this section: 1. How does the pattern of chromatic thresholds vary with model parameters? 2. Which part of the chromatic diagram should we sample to most efficiently estimate the model parameters?

I empirically estimated following parameters from MacAdam’s data: *Y* – luminance level of the *xyY* chromatic plane. *Y* scales all of the perceptual thresholds up and down, without changing their shapes. For model fitting, I additionally varied ***β*** – the proportion of each cone type. In Figure 2, I illustrated chromatic thresholds for four sets of ***β*** weights.

**Figure 2:**
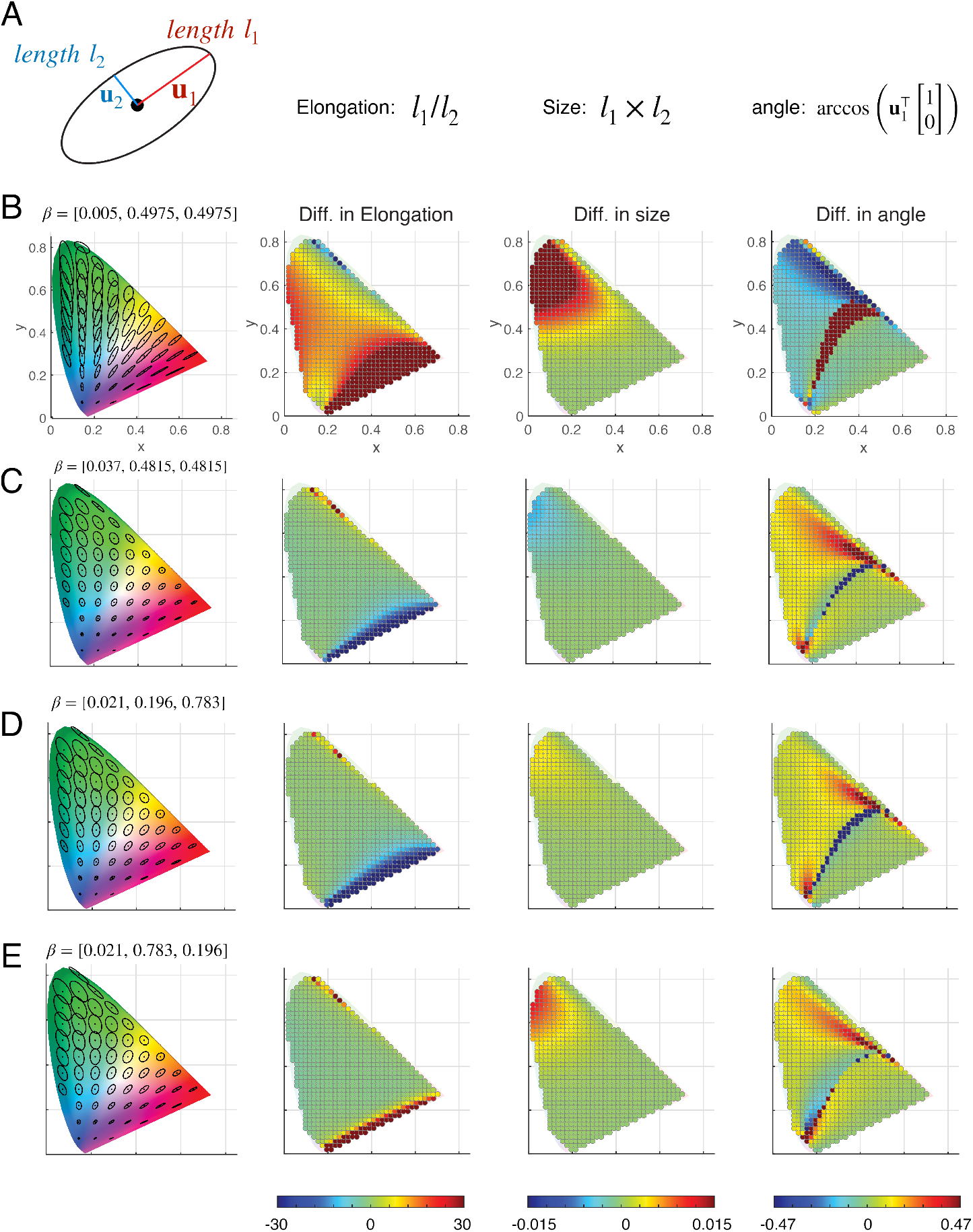
Parameter-induced change in chromatic thresholds. A. I summarized difference between two ellipses (two metric tensors) using three characteristics. Elongation is the ratio between the length of the major axis, and the length of the minor axis of an ellipse. Size is the product of the two lengths. Angle is the (cosine) angular difference between the major axis and the horizontal axis [1, 0]. Two pairs of ellipses are generally differ in these three aspects. B - E. Chromatic thresholds, as well as a break down summary of how each elliptical aspect vary (compared to those generated using ***β***^*∗*^) across the entire chromatic space for four different sets of ***β***.

The model predicted chromatic thresholds can be visualized as a 2-dimensional ellipse around a reference color (Figure 2A). First, I computed model-predicted thresholds for the optimal parameters found in Fonseca and Semengo 2016, ***β*** = ***β***^∗^ = [0.021, 0.4895, 0.4895]. Next, I varied ***β***, and compared the difference between the elliptical thresholds predicted using the new ***β***, versus those predicted using ***β***^∗^. In particular, I quantified the difference in three characteristics of an ellipse: the elongation of an ellipse (or eccentricity), the size, and the tilting angle. I used **u**_1_ and **u**_2_ to indicate the directions of the major and minor axes of an ellipse, and I used *l*_1_ and *l*_2_ to indicate the length of these two axes. A metric tensor (predicted chromatic threshold in the *xy* plane) is a positive semi-definite matrix, and it can be visualized as an ellipse, because it can be Eigen-decomposed as:

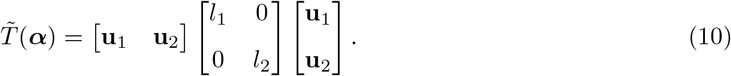

Elongation of an ellipse (commonly regarded as “eccentricity” in math) refers to the ratio between the length of its major and its minor axes, and in this case, 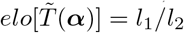. An ellipse is more elongated when it is narrow, and less elongated when it is round. The size of an ellipse is proportional to the square-root of the product between the length of its major and minor axes 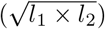, and without loss of generality, I computed a monotonic transform of the size, 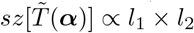. Combining 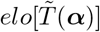 and 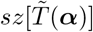. we can uniquely infer the set of *l*_1_ and *l*_2_. I use 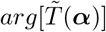 to indicate the angular difference between the major axis of the ellipse and the horizontal axis [1, 0], and the computation can be expressed as 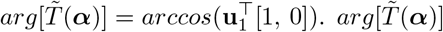 is 1 when the major axis of the ellipse is along the horizontal axis, and it is *π/*2 when the major axis is along the vertical axis.

To compare the difference in metric tensors predicted using two sets of ***β*** weights, first I plotted the difference in elongation between the two ellipses (at a single reference color, predicted using ***β***^∗^ and another pre-specified ***β***). The difference in elongation can be summarized using 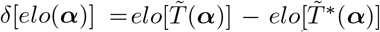. Similarly, I computed the difference in the size of the metric tensors as 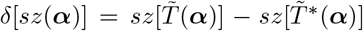, and the difference in how much each ellipse tilts as 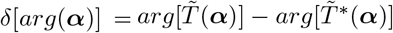.

In Figure 2B-E, I plotted *δ*[*elo*(***α***)], *δ*[*sz*(***α***)] and *δ*[*arg*(***α***)] for different reference colors (each reference corresponds to a cone response ***α***), and I visualized how much each characteristic varies across the entire chromatic diagram. It has been found that individual observers can have different proportions of *S, M* and *L* cone types within their retina, and the proportion of *M* versus *L* cones are especially diverse (Hofer et al. 2005; McMahon et al. 2008; Carroll, J. Neitz, and M. Neitz 2002; Deeb 2006). In Figure 2B, I decreased the proportion of *S* cones by 1.6% (relative to ***β***^∗^), and I increased it by 1.6% in Figure 2C. In Figure 2D, I decreased the proportion of *M* cone by about 30% relative to ***β***^∗^, and I increased it by about 30% in panel E. Even though the variation in the proportion of *M* and *L* cones are more drastic than the variation of *S* cones, from the first glance, the change in the three elliptical characteristics seem more sensitive to *S* cone variations. Additionally, we can make an observation that thresholds in some parts of the chromatic diagram seem to be more sensitive to ***β*** variations. This observation potentially leads to efficient sampling – to estimate model parameters, or to predict chromatic thresholds at un-sampled reference colors, we may only need to sample a handful of thresholds in the chromatic space.

To examine the possibility of more efficient sampling of chromatic thresholds, I further summarized *how much* the three elliptical aspects vary across the entire chromatic diagram and as a function of ***β***. First, I built a grid within the simplex of ***β*** weights anchored at *β*_*S*_, *β*_*M*_ and *β*_*L*_ (Figure 3A), and I fixed the luminance level *Y* for the rest of this analysis. I use ***β***_*i*_ to indicate the *i*^*th*^ set of ***β*** weights within the simplex. For each reference color, I quantified the difference in elliptical thresholds predicted using ***β***^∗^ and ***β***_*i*_ with the three elliptical attributes, and summed up the absolute value of the differences across all ***β***_*i*_ s within the simplex. The first three summary metrics (Figure 3B-D) can be expressed as the following:

**Figure 3:**
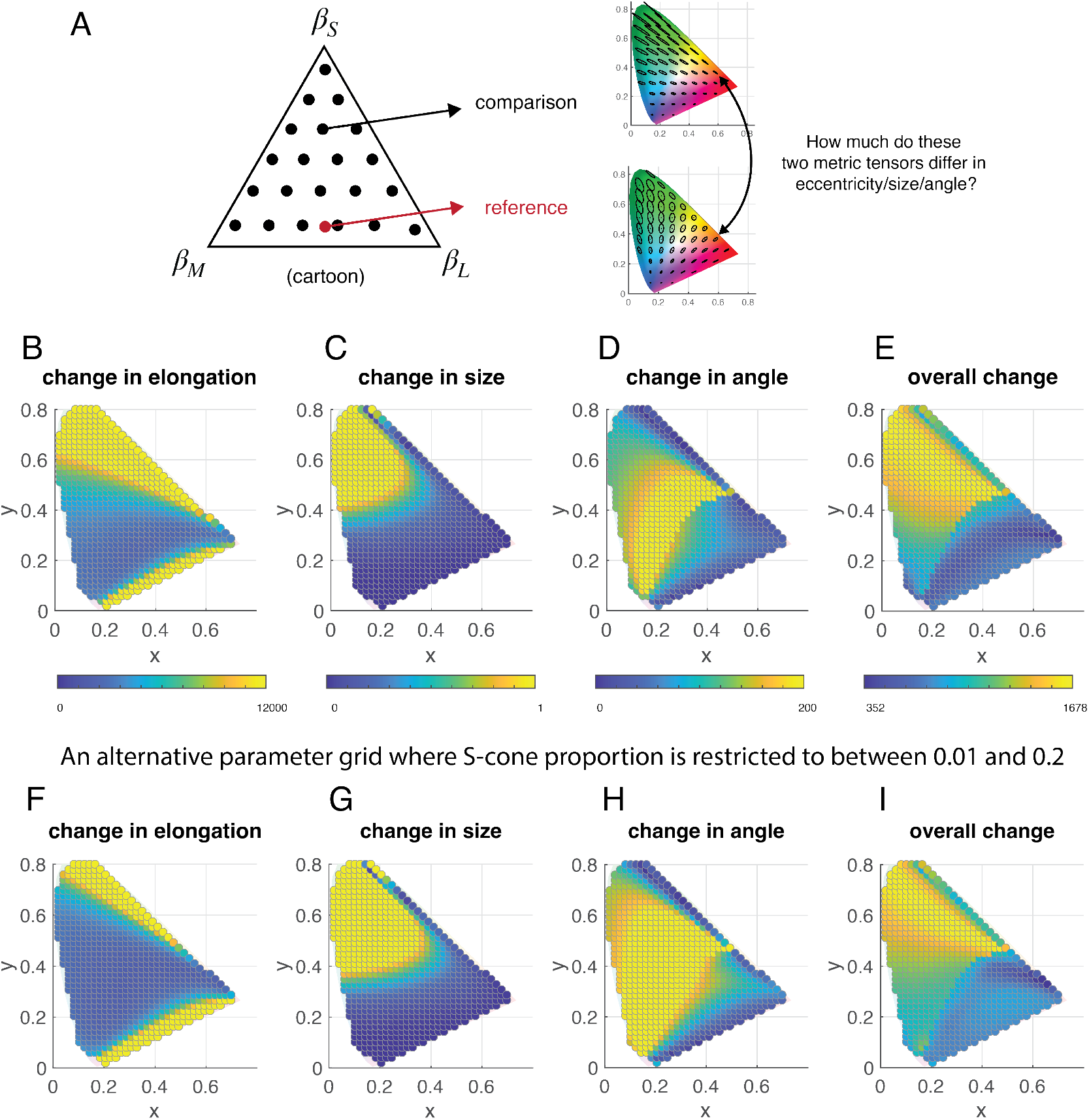
Efficient sampling in the chromatic space. A. Here, I illustrated a cartoon parameter grid – a simplex with the three *β* weights as anchor points. For each parameter set ***β***_*i*_ within the simplex, I generated a prediction for chromatic thresholds, and compared them to the thresholds generated using ***β***^*∗*^ – [0.021, 0.4895, 0.4895]. B-D. The comparison consists of three elliptical aspects (for predicted thresholds) at each reference point sampled in the chromatic space. The pattern of difference of all three elliptical aspects drastically differs. E. I combined the difference in all three elliptical aspects using Equation 14. F-I. Repeat the analyses in B-D but restricted the range of *β*_*S*_ to between 0.01 and 0.1.

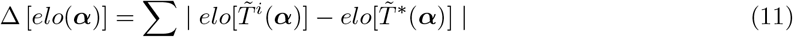

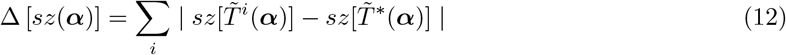

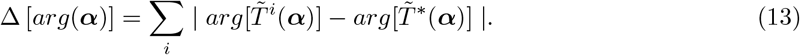

The ∆ maps are significantly different across the three elliptical characteristics. For example, the sizes of elliptical thresholds change the most at colors that correspond to high *y*-value, and low *x*-value in the chromatic diagram (or the green region). On the other hand, change in elliptical angles are the most significant within low- to mid-range *y* values (the blue-yellow region).

It would be simple and informative to summarize the overall elliptical changes (by combining the three elliptical aspects) using a single metric. The three elliptical characteristics have different units, so I cannot directly sum up the three ∆ maps. Alternatively, I ranked how much a metric tensor changes with respect to each elliptical attribute (across the chromatic diagram), and summed up the three ranking orders. Here is the detailed computation. For each metric tensor, I examined parameter-induced change in its elongation using ∆ [*elo*(***α***)]. I compared ∆ [*elo*(***α***)] across all reference colors (one for each ***α***), and ranked ∆ [*elo*(***α***)] across the chromatic diagram. The metric tensor that varies the least in elongation has a rank order 1, and the metric tensor with the most variation has a rank order *N*, and *N* is the total number of metric tensors. In a similar manner, I computed the rank order for ∆ [*sz*(***α***)] and ∆ [*arg*(***α***)]. To visualize to overall parameter-induced variations in metric tensors, I plotted the following quantification computed for reference colors across the chromatic space:

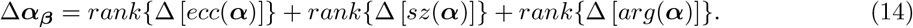

Overall, the metric tensors that situate at mid- to high-range of *y* values and mid-range of *x* values (the green-yellow region) are more informative about model parameters than others.

To testify chromatic thresholds measured in the said region are more informative about model parameters, I conducted an additional analysis. First, I chose three regions on the chromatic diagram according to Figure 3I, region 1 - 3 (Figure 4A). Chromatic thresholds in region 1 varies a lot with change in ***β*** (the most informative region), and less so in region 2, and the least in region 3 (the least informative region). Within each region, I randomly sampled reference colors, at which I used ***β***^∗^ = [0.021, 0.4895, 0.4895] to generate metric tensor predictions. In real psychophysics experiments, we measure perceptual threshold along one image perturbation direction at a time. To generate simulations that assimilate psychophysical threshold data, I sampled the length of 4 or 8 lines (along randomly chosen directions) within each metric tensor (Figure 4B). Then using these line samples, I tried to estimate the model parameters back (see Methods). In Figure 4C - E, I plotted parameter recovery for *β*_*S*_ with increasing number of reference colors sampled within the 3 regions. I generated 4 line samples from each metric tensor, and used them as data for parameter recovery. In Figure 4F - H, I plotted the same parameter recovery analysis but with 8 line samples generated for each metric tensor. In general, parameter recovery from region 1 is a lot more reliable than those from the two other regions. For example, with 8 line samples, using as few as 2 reference colors, we are likely to recover the proportion of *S* cones. The efficiency in parameter recovery suggested that potentially, we can only sample a handful of chromatic thresholds to predict the entire pattern of thresholds for any point on the chromatic diagram.

**Figure 4:**
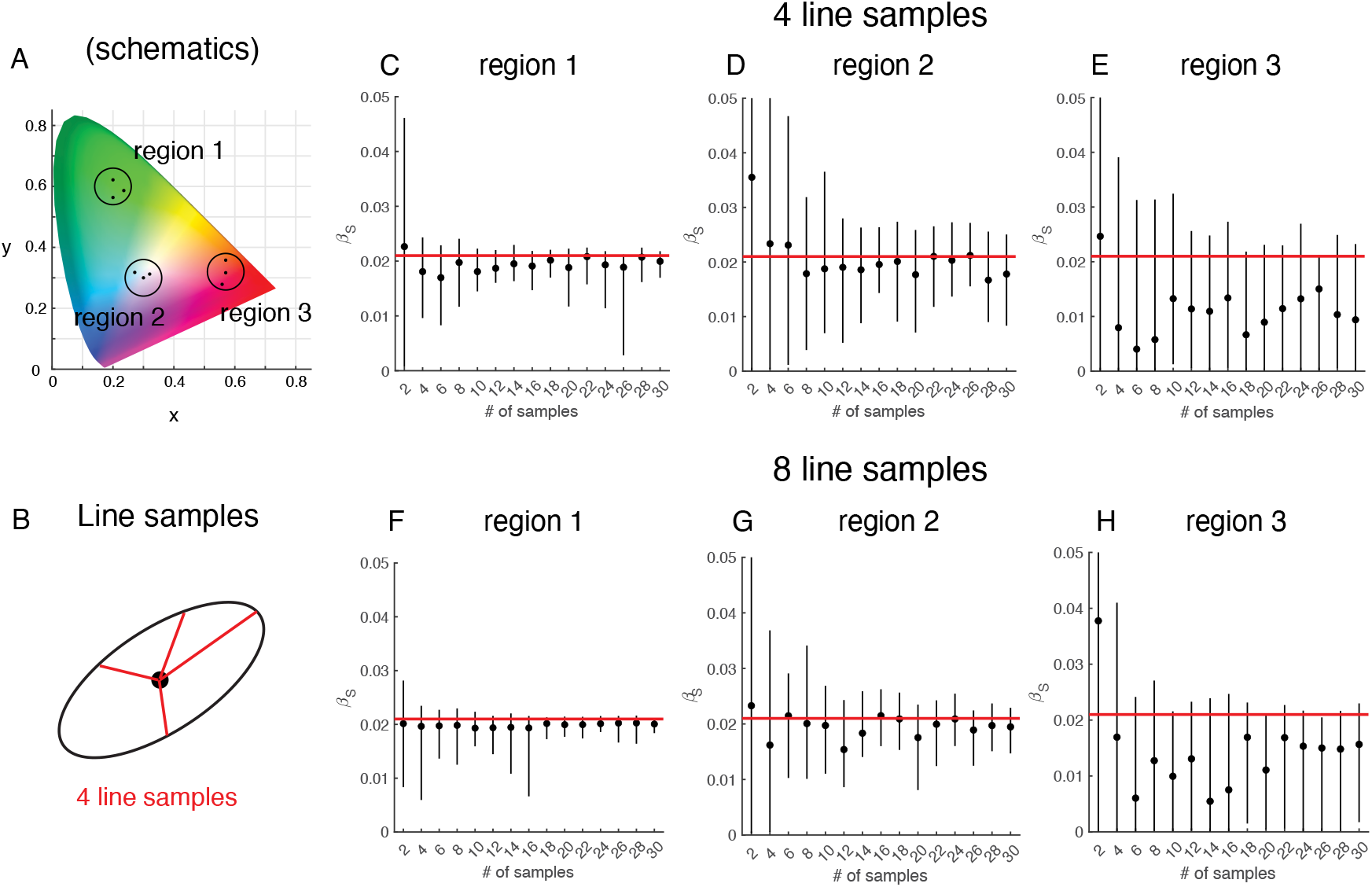
Parameter recovery in different regions of the chromatic diagram. A. Schematics. I picked three regions on the chromatic diagram. Region 1 corresponds to chromatic thresholds that vary a lot with changing ***β*** weights (informative region). Region 2 is less informative, and region 3 is the least informative. B. For parameter recovery analysis, I randomly sampled reference colors within each region. For each reference color, I used the model (linear neural model with Poisson noise) to generate metric tensor predictions. I randomly sampled either 4 or 8 directions from each predicted metric tensor and I added Gaussian noise (*σ*= 0.1) to each line sample. Each line sample assimilates a threshold measurement in psychophysics data. C-H. I varied the total number of reference colors that I sampled from each region (such process is repeated 100 times to generate error bars), and estimated the proportion of *S* cones (*β*_*S*_). I plotted the input *β*_*S*_ = 0.021 (the red line) versus the predicted *β*_*S*_. In general, sampling in region 1 is more informative about *β*_*S*_ than sampling in the other two regions. Error bars represent the central 40% quartile range.

### 2.3 Improved Prediction of chromatic thresholds using alternative noise models

In this section, I first examined how much the Poisson noise assumption contributes to the predicted chromatic thresholds, then I analyzed alternative noise models.

#### The contribution of Poisson noise to predicting chromatic thresholds

In previous sections, I re-produced the chromatic threshold predictions using a linear model paired with Poisson noise. The predicted thresholds (in the *LMS* space) were further passed through additional nonlinear transforms to be visualized in the *xy* chromatic diagram. It is important to know which of the three computations the linear cone response, Poisson noise, or the non-linear mapping between different color spaces, contributes the most to the chromatic threshold predictions? To tease these elements apart, I did an ablation analysis: I removed the Poisson noise from the model, and re-examined the threshold predictions. The neural model becomes deterministic without the Poisson noise. Psychophysical experimenters do not typically consider deterministic models, because these models cannot explain observers’ stochastic behavior in a threshold experiment. Nevertheless, deterministic models can still be used to make predictions to *relative* thresholds measured across stimuli. In the case of chromatic thresholds, instead of computing Fisher thresholds, (local) distance traversed in the neural response space can be summarized using the Jacobian of mean neural response functions (J. Zhou and Chun 2022) (see Methods). With linear neural response ***α***, the metric tensor in the *LMS* space has the form of the identity matrix:

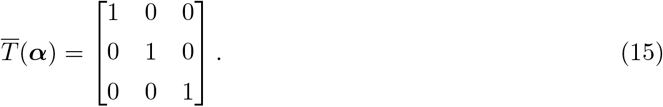

I further transformed this metric tensor to the *xy* chromatic diagram following Equation 8. For details of this computation, see J. Zhou and Chun 2022 and the Method section.

One would have expected that the Poisson noise could be crucial for achieving good chromatic threshold predictions, because it is the only non-linear component in the neural model. Since MacAdam ellipses are diverse in shapes and sizes, and a linear (overall) model are not sophisticated enough to produce the intricate threshold patterns. Surprisingly, the deterministic model better captures the MacAdam ellipses than the stochastic model (Figure 5AB). This indicates that the Poisson noise is not only inconsequential in predicting chromatic thresholds, it also could have hindered the accuracy of the predictions. Because the linear part of the neural model would not contribute to the diverse shapes of the MacAdam ellipses J. Zhou and Chun 2022, the only non-linear component left is the non-linear transform from the *XY Z* space to the *xyY* chromatic diagram.

**Figure 5:**
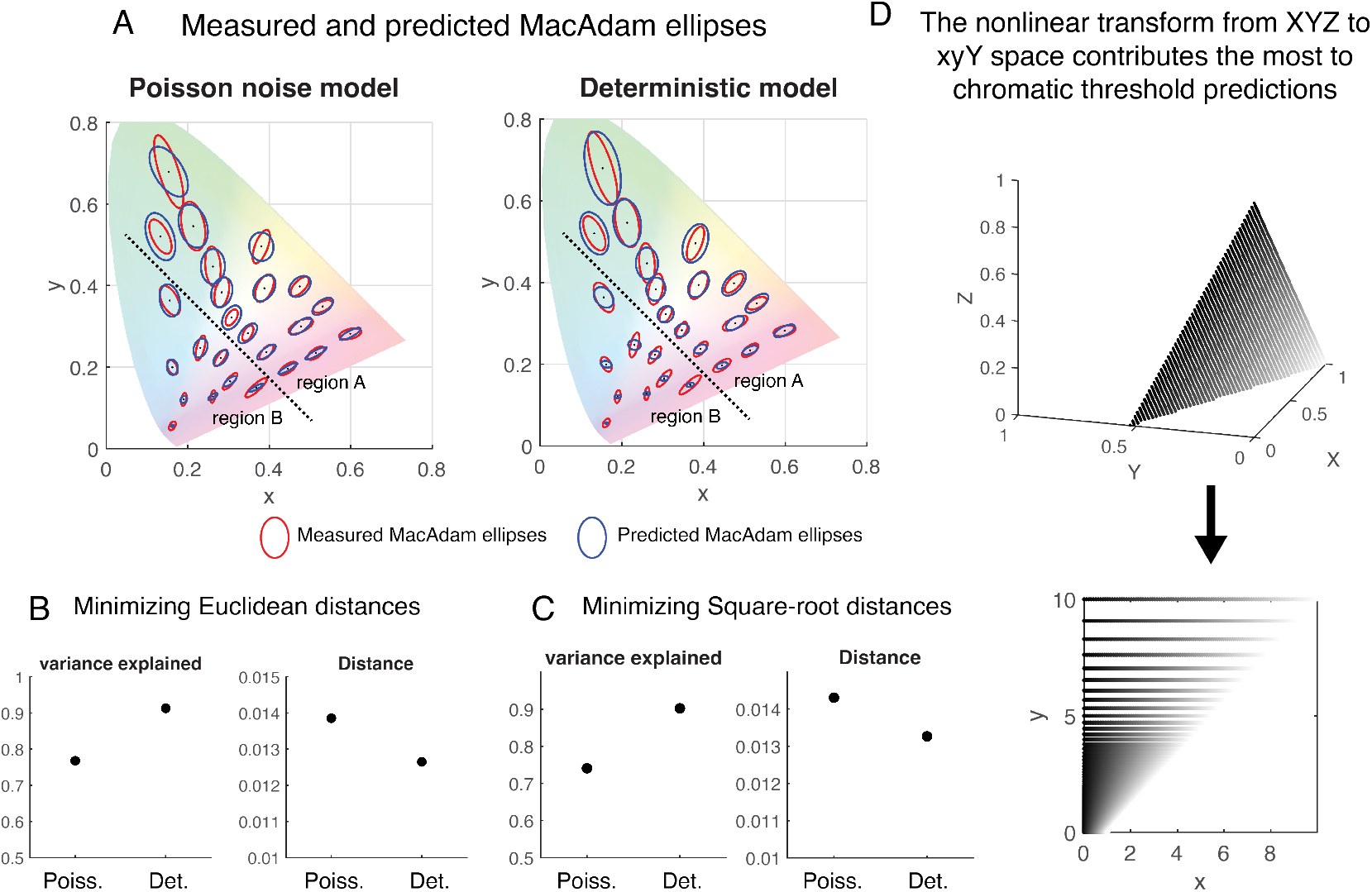
Assessing to what extent Poisson noise contributes to predicting chromatic thresholds. A. Both the deterministic model, and the linear model assuming Poisson noise reasonably captured MacAdam’s measurements (red). B. Fitting both models by minimizing the Euclidean distance between the measured and the predicted metric tensors. The deterministic model achieved higher variance explained (0.91), compared to the Poisson model (0.77). I used a second metric to summarize the quality of model fits. This metric sums up the Frobenius distances between the measured and the predicted metric tensors. Smaller distance means better model predictions. C. The same analyses as panel B, but square-root distance was used (instead of Euclidean distance) during optimization. D. The nonlinear transform (luminance normalization) from *XY Z* color space to *xyY* space is crucial for capturing the diverse shapes and sizes of the MacAdam ellipses.

In Figure 5D, I used a simulation to illustrate how the nonlinear transform gave rise to the chromatic threshold pattern. I evenly sampled on a hyperplane with reduced range of *Y* values in the *XY Z* color space. This is because to account for experimental data, I sampled reference colors in the *xy* space with no luminance variations. Once these reference colors are projected to the *XY Z* space, they lay on a hyperplane that barely varies along the *Y* axis. In this simulation, I mapped the simulated hyperplane in the *XY Z* space to the *xy* space via Equation 3. The output in the *xy* plane has varied distance structures that resembles what we observed in the pattern of chromatic thresholds. The distance between sample points becomes larger with increasing values of *x* and *y*, and closer near the origin, and metric tensors like those measured in chromatic thresholds reflect the underlying distance structures. Equation 3 is essentially a process of luminance gain control. It is perplexing that luminance gain control has been playing the most crucial role in predicting the pattern of chromatic thresholds.

#### Predicting chromatic thresholds using alternative noise models

Since Poisson noise model hinders the chromatic threshold predictions, what would be a better noise model, if any, that improves the perceptual predictions? To answer this question, I examined three additional types of noise in this section.

In Figure 6A, I showed re-produced Poisson noise prediction to MacAdam ellipses, which closely resembles the predictions in Fonseca and Semengo 2016. The first additional noise model that I examined was equi-variant Gaussian noise. This model contains one additional parameter *σ* compared to the Poisson noise model, and *σ* is the Gaussian standard deviation that is assumed the same for all three cone types. Under this assumption, the predicted thresholds in the *LMS* response space has the following expression:

**Figure 6:**
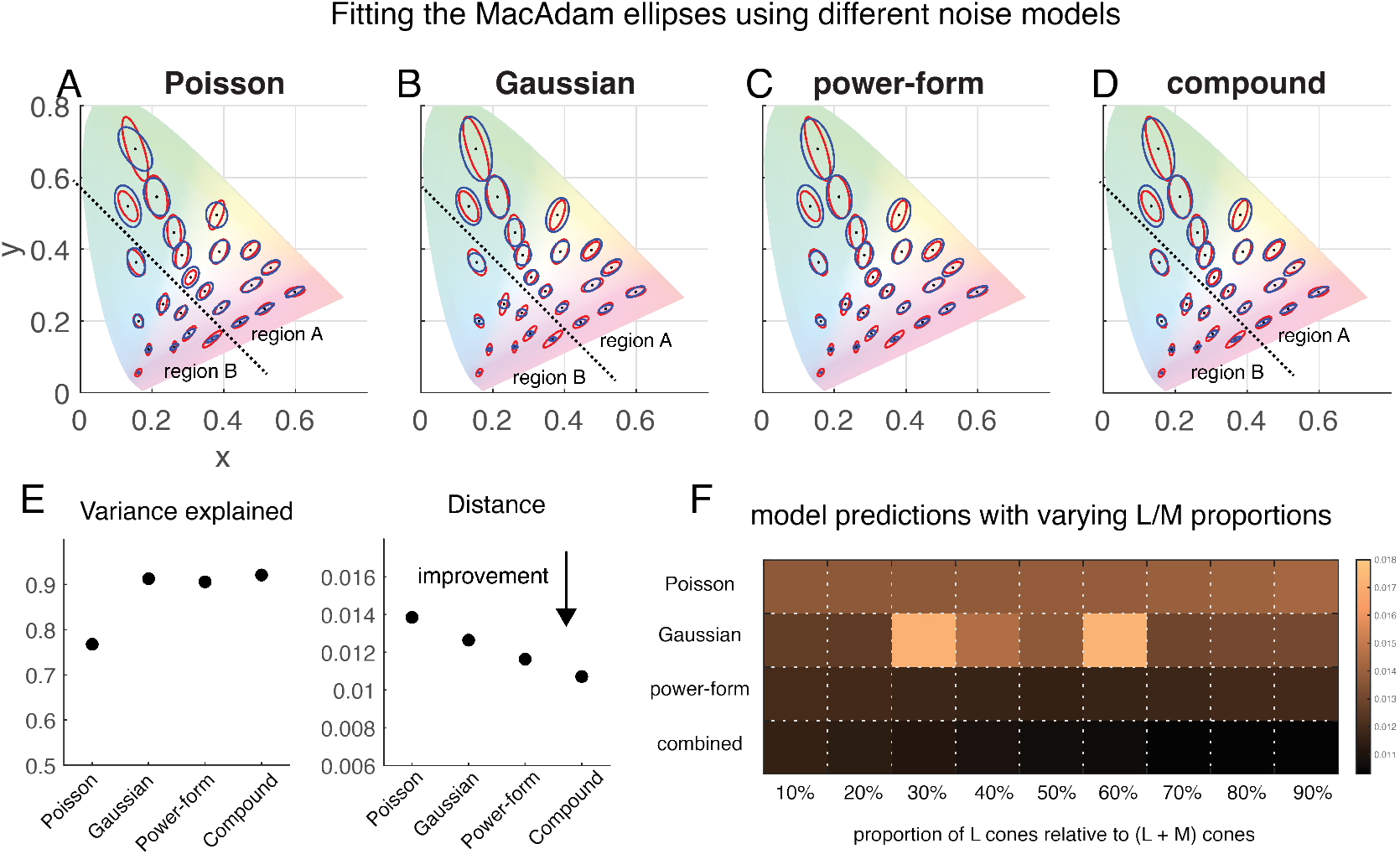
Chromatic thresholds predicted using different noise models. A-D. Chromatic thresholds predicted using independent Poisson, Gaussian, power-form noise model, and compound noise model. The red ellipses are re-produced MacAdam chromatic thresholds, and the blue ellipses are model predictions. E. The Euclidean distance between measured and predicted chromatic thresholds for each noise model, the lesser distance is better. F. The same distance metric for different proportions of L versus M cones. In panel A-E, *β*_*S*_ was empirically estimated assuming *β*_*M*_ = *β*_*L*_. Here, I varied the proportion between *L* and *M* cones, and for each proportion, I re-estimated *β*_*S*_. I plotted the Euclidean distance between each model’s prediction and measured chromatic thresholds. Smaller distance (darker shades) means better model fits.

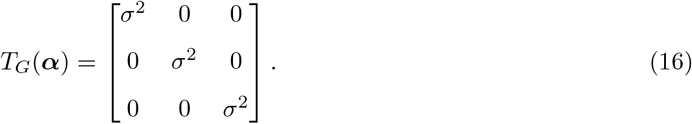

This model is equivalent to the deterministic model in the sense that both *σ* and the luminance level *Y* control the overall size of all chromatic threshold ellipses, so there is a trade-off between the two parameters, and neither parameter is uniquely identifiable. As a consequence, the predicted chromatic thresholds using this model (Figure 6A) is also equivalent to the deterministic model’s predictions.

The second alternative noise model I examined has a relatively flexible form of standard deviation (Figure 6C). I assumed that the noise standard deviation for each cone type is a power function of its mean response. For example, for *S* cone, the noise standard deviation *σ*_*S*_ is assumed to be 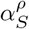, and *ρ* is a non-negative exponent. If *ρ* is 0, this model is the same as the equi-variant Gaussian model. If *ρ* is 2, the model is the same as the Poisson noise model. When *ρ* is 1, the model is similar to the super-Poisson noise that has been observed in neural responses in visual cortices (J. Zhou, Duong, and E. P. Simoncelli 2022; Lin et al. 2015; Goris, Movshon, and E. Simoncelli 2014). Additionally, *ρ* can take any other non-negative values. For this model, the additional parameter that we need to estimate is *ρ*, and *ρ* is assumed to be the same for all three cone classes.

The third alternative noise model results from an observation that comparing between the Gaussian and the Poisson model predictions, the Gaussian model seemed to make veridical predictions in region A (Figure 6AB), whereas the Poisson model seems to make better predictions in region B. So potentially, by combining the two noise models, an improved prediction can be achieved in both regions (Figure 6C). Here, I examined a combined model that was previously suggested for other modalities (J. Zhou, Duong, and E. P. Simoncelli 2022). I used *S* cone as an example to illustrate this noise model. For *S* cones, I assumed the noise standard deviation 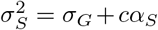. As in the case of Gaussian noise, *σ*_*G*_ is a constant to be estimated. *c* is an additional scaling constant that is non-negative, and *α*_*S*_ is the mean *S* cone response. Threshold metric tensor derived for this noise form has the following expression:

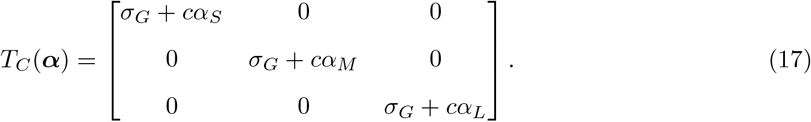

Each of these three models, in addition to the Poisson noise model were fit to empirically measured MacAdam ellipses. For each model, I varied the luminance level *Y*, the proportion of *S* cones *β*_*S*_, and additional parameters that concern neuronal noise (see Methods). All three additional noise models improved upon the Poisson noise model, and I quantified the model fits using two different metrics. First, I quantified how much variance in the data is explained by each model (see Methods). All three additional noise models managed to account for more 90% of the data variance, and showed improvements upon the Poisson model predictions (Figure 6EF). Second, I quantified the goodness of fits using summed distance between the empirically measured ellipses and the model predictions. Let *d* [*T*_*G*_(***α***), 𝒯 (***α***)] be some distance metric between the model-predicted metric tensor (with Gaussian noise as an example) *T*_*G*_(***α***), and the empirically measured metric tensor at ***α***, 𝒯 (***α***):

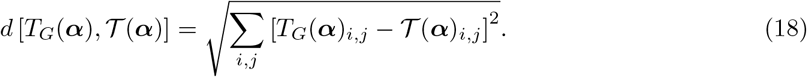

This is equivalent to a sum of the Frobenius norm between the matrix of measured metric tensor, and the matrix of the predicted metric tensor. The shorter distance means a better model prediction. Using this criterion, the noise model that combines the Gaussian and the Poisson noise out-performed other models’ predictions.

#### 2.4 Model-predicted chromatic thresholds across retinal eccentricities

Being able to predict chromatic thresholds can greatly enhance our understanding of many chromatic properties of perception. In this section, I illustrate an example application. Using a linear cone model combined with the compound noise model (as described in previous section), I made predictions to how chromatic thresholds vary across retinal eccentricities.

I extracted retinal data from Weinrich et al. 2017, which summarized that across retinal eccentricities, how the overall cone density and the proportion of *S* cones decay. The retinal data were averaged across 3 young (6 years old) primates (*Macaca fascicularis*) measured using epifluorescence imaging. I fitted an exponential function with three parameters to capture the decaying shape in both (M + L) cone densities (Figure 7A), and in S cone densities (Figure 7B). The exponential function has the following form: *ae*^−*bx*^ + *c*, and *x* is the retinal eccentricity measured in millimeters. *a, b, c* are empirically estimated parameters. In Figure 7C, I illustrated the total cone density and the *S* cone density normalized to the cone density at *ecc*. = 2*mm* (which is set to 1). In Figure 7D, I showed the *S* cone density relative to the total cone density at each eccentricity (so that the total cone density is assumed 1 for all eccentricities).

**Figure 7:**
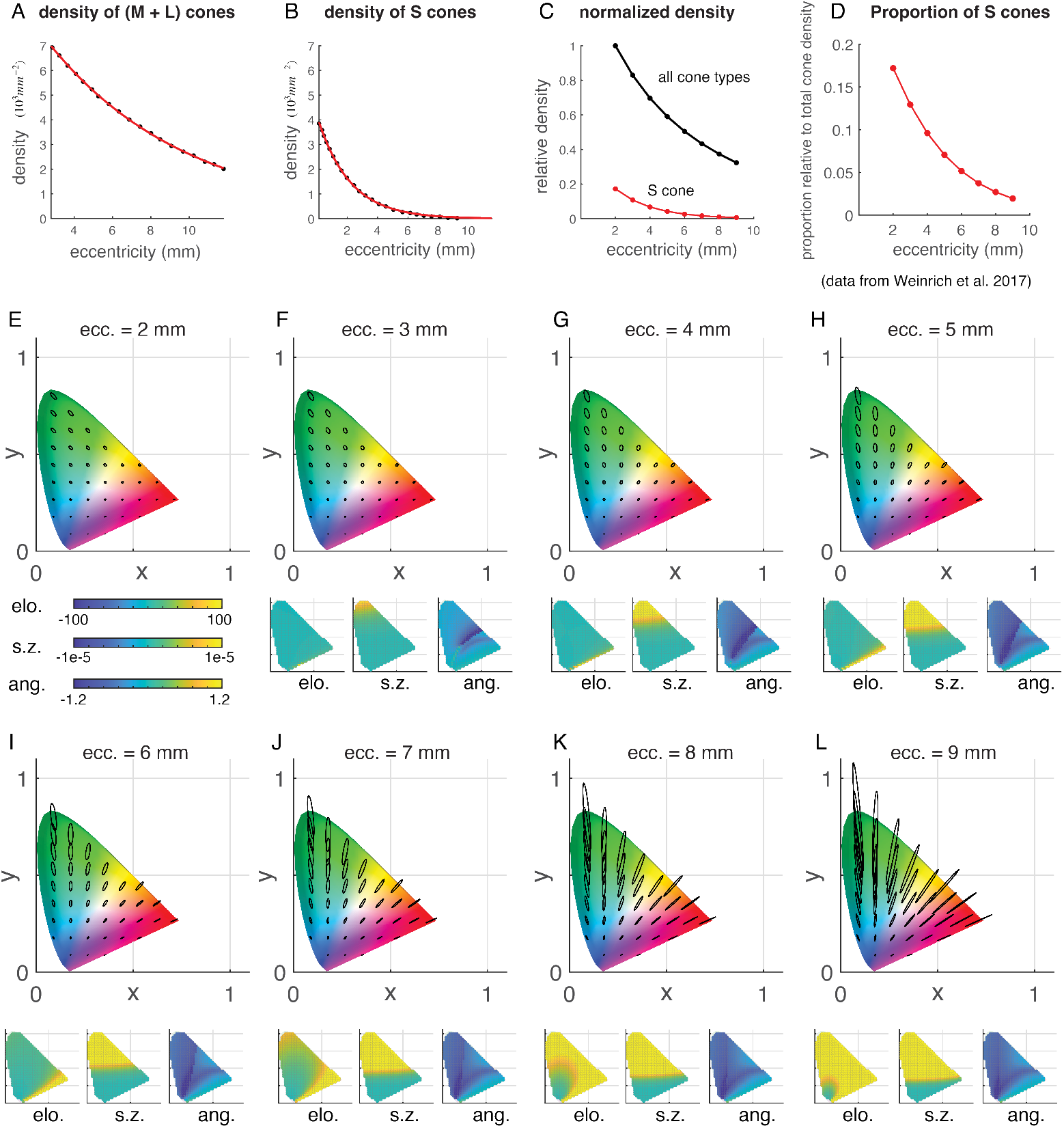
Predicting chromatic thresholds varied across retinal eccentricities. A. Cone density measured for (M + L) cones in Weinrich et al. 2017 (black dots), and the red curve is the exponential fit to the data with 3 varying parameters. B. Data and exponential fit to the S cone density in primate retinae (data averaged across 3 primate retinae). C. Normalized overall cone density and S cone density, assuming that the overall cone density at 2 mm reitnal eccentricity is 1. D. Proportion of S cones relative to overall cone density (normalized to 1) for eccentricities rangin from 2 to 9 mm. E - L. Predicted chromatic thresholds across retinal eccentricities using the compound noise model. Additionally, I summarized the difference between the pattern of chromatic thresholds at each eccentricity, and those at 2 *mm* eccentricity using the three elliptical characteristics.

In Figure 7E-l, I summarized the predicted chromatic thresholds across retinal eccentricities using the compound noise model. I plotted the predicted chromatic thresholds (sparsely sampled) at different eccentricities, and illustrated how these predictions differ from the chromatic thresholds predicted at *ecc*. = 2*mm*. I summarized these differences using the three elliptical characteristics (densely sampled). Across all eccentricities, perceptual thresholds are the largest in the green region in the chromatic diagram (low *x* and high *y* values), and are the smallest in the blue/purple region (low *x* and low *y* values). Both size and shape of the chromatic thresholds significantly vary across eccentricities. There are two contributing factor to the variations, the total number of cones, and the proportion of S cones. The size of chromatic thresholds generally decreases with total number of cones, and less cones (as in periphery) leads to less precise color encoding. The total number of encoding cones do not affect the shape of the chromatic thresholds. So the variation in elongation and angle (or shape in general) of the chromatic thresholds are attributed to the variation in the proportion of *S* cones. Overall, perceptual thresholds tend to vary the most in the green regions in the chromatic diagram, whereas change in the angle of the elliptical representations seem most significant along the blue-yellow region in the chromatic space.

Previous literature (e.g. Hansen, Pracejus, and Gegenfurtner 2009) suggested that (1) color vision is the best in fovea and declines in the periphery (see Johnson 1986 for a summary); (2) Sensitivity to red-green color variations declines more steeply toward the periphery than sensitivity to blue-yellow color variations (e.g.Mullen, Sakurai, and Chu 2005); (3) sensitivity to some hues (e.g. green) declines faster than others (e.g. red) (Boring 1942; Stromeyer, Lee, and Esker 1992; Abramov, Gordon, and Chan 1991). To some extent, all of these observations are reflected in Figure 7, and in future work, the model predictions across retinal eccentricities will be tested against human data, and more quantitatively compared to existing color vision evidence in human periphery.

## Discussion

In this work, first, I studied two aspects of chromatic perception using an existing cone model. I quantified how the pattern of chromatic thresholds varies when the proportion of three cone types varies. This analysis led to efficient sampling of chromatic thresholds. Second, I analyzed to what extent the assumption of Poisson noise in the existing neural model contributes to the chromatic threshold predictions. Surprisingly, Poisson noise is inconsequential to the prediction, and it is the nonlinear projection of the perceptual thresholds from *XY Z* to *xyY* color space (a process of luminance gain control) that plays the major role. Additionally, I analyzed three alternative noise models, each of which achieved improved chromatic threshold predictions, compared to the Poisson noise model. Last, I showed an example of how model-predicted chromatic thresholds can be used to guide new data collection. I examined how cone density and the proportion of *S* cones vary across retinal eccentricities, and these variations lead to drastically different chromatic threshold patterns across retinal eccentricities.

The measurements of chromatic threshold do not lead to unique identification of an underlying neural model. A vast psychophysics literature has examined this identifiability issue with one-dimensional stimulus perturbations (e.g. Katkov, Tsodyks, and Sagi 2007; García-Pérez and Alcalá-Quintana 2009; A. J. Solomon 2007; F. Kingdom 2016). For example, Weber’s law, a common form of perceptual discriminability that was observed in many stimulus domains, can be implemented by many different internal representations (J. Zhou, Duong, and E. P. Simoncelli 2022). This is because if we observe Equation 1, both the mean and the variance of the internal representation contribute to perceptual discriminability, and from observing discriminability alone, we cannot uniquely determine either the representational mean or variance. This identifiability issue persists in the case of two-dimensional (or *n*-dimensional) perceptual discriminability. Many different internal or neural representations can gie rise to the same prediction of chromatic threshold pattern. In Figure 8, I showed an example where a pattern of chromatic thresholds can be generated from two seemingly different neural representations – a linear neural model that assumes equi-variant Gaussian noise, and a quadratic neural model that is paired with Poisson noise.

**Figure 8:**
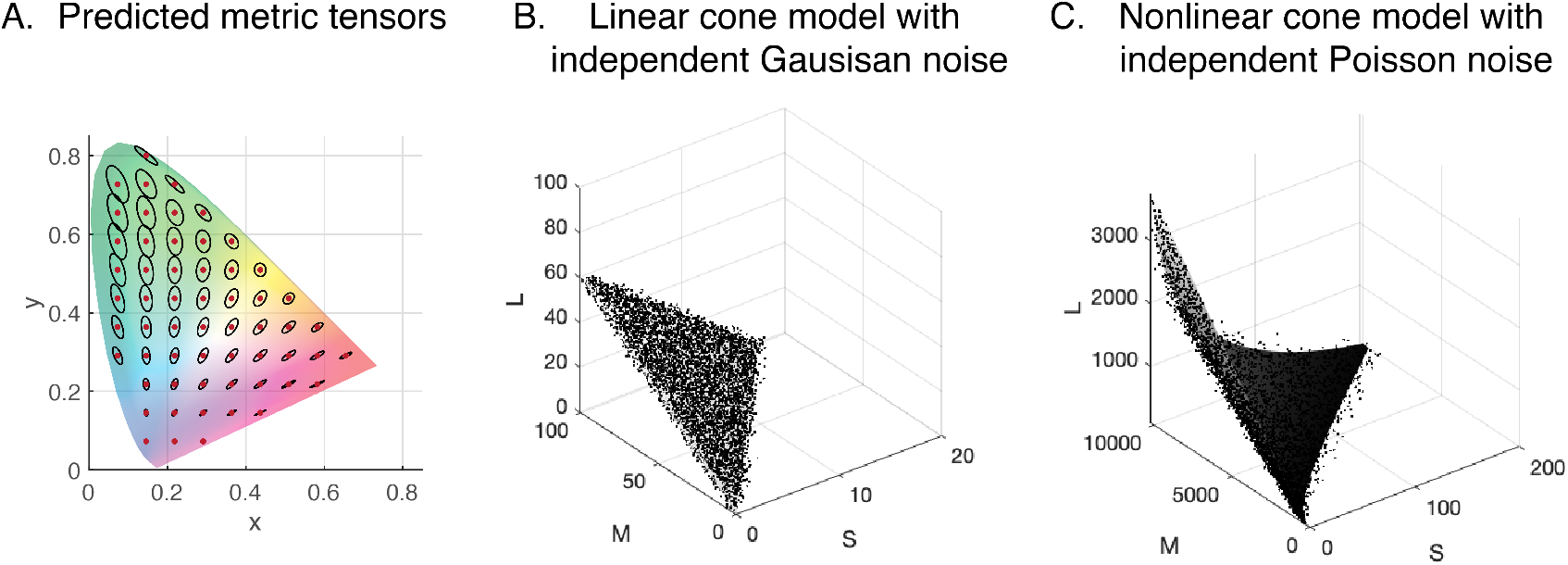
Identifiability issue in two-dimensional perceptual discriminability. We cannot uniquely identify an internal representation (or a neural representation) from measurements of discriminability alone. A. The same pattern of chromatic thresholds as Figure 1B. B. A linear neural model that is paired with the Gaussian noise (equi-variance) that give rise to the chromatic threshold predictions shown in panel A. C. A nonlinear cone model that is paired with independent Poisson noise that give rise to the same pattern of chromatic threshold predictions.

An alternative model that predicts metric tensors in the DKL color space was proposed in Duinkharjav et al. 2009. In Duinkharjav et al. 2009, the authors empirically measured chromatic thresholds in the DKL color space, and used a radial basis function neural network (with a sigmoidal output layer) to approximate (via regression) the measured data. The measured thresholds, like in the previous literature, exhibit following properties: (1) the authors observed unequal thresholds with different reference colors even at the same retinal eccentricity; (2) discriminability decreases in general as retinal eccentricity increases; (3) inter-subject variations were observed. Even though directly comparing the neural model that predicts chromatic thresholds in the *xy* space, to the neural network model developed to capture metric tensors in the DKL space (Duinkharjav et al. 2009) is out of the scope of this paper, these computational approaches to model metric tensors in color present a unifying effort to develop colored displays that are customarily calibrated for each individual.

## Methods

### Computing threshold predictions using a deterministic neural model

Traditionally, stochastic neuronal models are used to predict perceptual thresholds, assuming that observers’ stochastic perception (e.g. observers’ varied responses to discriminating between a pair of stimuli) comes from stochastic underlying representations. Without the noise assumption, there is no uncertainty in observers’ perception. However, in comparative threshold studies – studies that compare perceptual thresholds across different stimulus perturbations, deterministic models can still be useful.

A single reference color in the chromatic diagram (the *xyY* color space) can be perturbed along any direction in the two-dimensional *xy* plane, assuming that all such perturbations are unit-length vectors, and all perturbation to the reference forms a unit circle. Previous experimental evidence supported that perceptual discriminability reflect the extent of change in underlying neuronal responses, and in particular, the extent of change is consistent with a summary using the *L*2 norm (Poirson and Wandell 1990; Knoblauch and Maloney 1996; J. Zhou and Chun 2022). The consequence of the L2 norm is that perceptual discriminability (hence thresholds) at a reference color exhibit an elliptical shape. The elliptical shape is the visualization of a positive definite matrix, which is called a metric tensor, and can be derived from a neural model. For stochastic neural models, we can derive metric tensors using Fisher information, and for deterministic models, we can also derive metric tensors, using the Jacobian of the model. More detailed derivation of metric tensors from directional derivative of the model and model Jacobians, see J. Zhou and Chun 2022.

Here is a brief summary. I use ***µ***(**s**) to denote the deterministic neuronal transform ***µ*** as a function of *p*-dimensional stimulus **s**, and the output of ***µ***(**s**) is an *n*-dimensional neuronal response. To summarize how the neuronal response ***µ***(**s**) varies when stimulus **s** is perturbed along the direction (assuming ||***ϵ***|| = 1 for simplicity), we can calculate the directional derivative of ***µ***(**s**) with respect to **s** along the ***ϵ*** direction. The directional derivative **d**_***ϵ***_(**s**) can be expressed as the following:

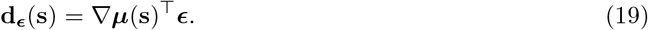

Since discriminability reflects *how much* the neuronal response changes, to make predictions to discriminability, we need to calculate ||**d**_***ϵ***_(**s**) ||, the *L*2 norm of the directional derivative, which has the following expression:

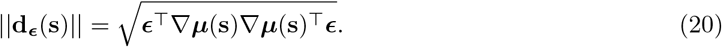

∇***µ***(**s**) ∇ ***µ***(**s**)^⊤^ is the metric tensors. It is a positive definite matrix, and it can be visualize as an ellipse in the case analyzed in the main text, and it totally determines how each direction of stimulus perturbation ***ϵ*** is transformed in the neuronal representation. Since thresholds are the inverse of perceptual discriminability, and by computing the inverse of this metric tensor, we obtained the metric tensor that represents perceptual thresholds.

### Computing Fisher information

Metric tensors for a stochastic neural model can be similarly derived, by replacing the metric tensors for the deterministic model with Fisher information. Fisher information is difficult to compute for general distributions, and there is no guarantee of a closed-form expression. So in practice, we often instead compute a lower bound on Fisher information (Stein, Mezghani, and Nossek 2014; J. Zhou, Duong, and E. P. Simoncelli 2022), and this lower bound is exact when the underlying neuronal response distribution is within the exponential class, and has linear sufficient statistics (Kfashan et al. 2021). In other words, this lower bound is exact for many commonly used distributions (e.g. Poisson and binomial). Essentially, in the deterministic model, the metric tensors are completely determined by the Jacobian of the (averaged) neuronal responses, or the change in the population response. In the stochastic case, the metric tensors are determined by the change in the population responses, as well as the covariance of the population response, as expressed in Equation 2.

### Transforming metric tensors

To understand transformations of metric tensors, I started by analyzing a deterministic model, before doing an analogous computation for stochastic neural models. Assuming a deterministic neural model ***µ***(**s**), if we transform this neural response via a vector function **g**[***µ***(**s**)], the Jacobian of the transformed function has the form ∇**g** ∇***µ***(**s**) because of the chain rule. Because metric tensors summarized the extent of stimulus-perturbed response in the transformed **g** space, we can correspondingly compute the metric tensor as previously described, which as a form ∇**g** ∇***µ***(**s**) ∇ ***µ***(**s**)^⊤^ ∇**g**^⊤^. The stochastic computation is very similar, we can replace ***µ***(**s**) by the computation of the lower bound on Fisher information.

### Generating line samples

In simulations for Figure 4, I generated line samples from a metric tensor to recover the model parameters. A line sample can be described as the following. Given a metric tensor 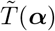, to generate experimental data that is like perceptual thresholds, I generated some random two-dimensional unit vector ***ϵ***, and computed

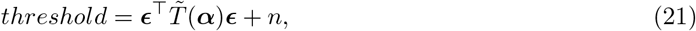

such that *n* is a Gaussian random variable representing experimental noise.

### Model Fitting

I fitted models to MacAdam’s data using two objective functions. Each model predicts a metric tensor (a 2 ×2 positive definite matrix) at each reference color, and MacAdam’s ellipses consist of an ellipse measured at each reference color. First, from each MacAdam’s ellipse, I deduced a (measured) metric tensor. This is because knowing the shape of an ellipse, we know the directions and lengths of its major and minor axes, from which we can reconstruct the metric tensor using Equation 10.

After reconstructing the measured metric tensors, for each pair of measured and predicted metric tensors, I compared their distances, and then I summed up the distances across all pairs of metric tensors. I used two different metrics to quantify the distance. First, I used Frobenius norm to quantify distances between a measured metric tensor 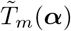, and the corresponding prediction 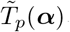. The forbenius norm can be expressed as:

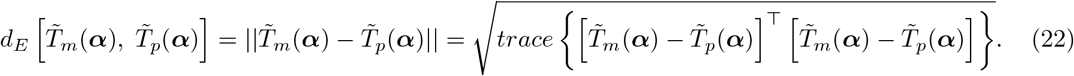

Because the space of symmetric positive definite matrices is not an Euclidean space, and some other choices of quantifying distances between such matrices could be more natural. A an alternative metric, I minimize (the sum of) the Frobenius norm of the difference between the square-root of the metric tensors, expressed as:

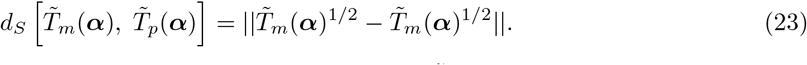

Here is how to compute matrix square root. Suppose, for example, 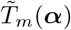 can be eigen-decomposed as 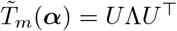, the matrix square-root 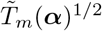 can be found via 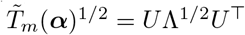. For additional metrics that can be used to compare between two covariance matrices (or two metric tensors), see Dryden, Koloydenko, and D. Zhou 2009.

### Quantifying model fits

In Results, I used two ways to quantify how well a model fits the data. The first quantification is the variance explained. Each model predicts a set of metric tensors (2× 2 symmetric positive definite matrices), and the measured MacAdam ellipses can also be represented as a set of metric tensors. To calculate variance explained, I used 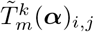 to denote an entry of the *k*^*th*^ metric tensor 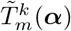, and do the following calculation:

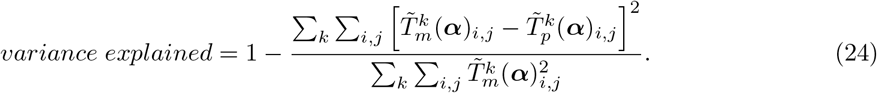

The second quantification I used was the sum of the Frobenius norm between the measured and the predicted metric tensors.

## Acknowledgement

I thank Eero Simoncelli and Zhong-Lin Lu for inspiring conversations and insightful questions.

## Notes

### Competing Interest Statement

The authors have declared no competing interest.

